# Development of adaptive motor control for tactile navigation

**DOI:** 10.1101/762443

**Authors:** Alireza Azarfar, Tansu Celikel

## Abstract

Navigation is a result of complex sensorimotor computation which requires integration of sensory information in allocentric and egocentric coordinates as the brain computes a motor plan to drive navigation. In this active sensing process, motor commands are adaptively regulated based on prior sensory information. In the darkness, rodents commonly rely on their tactile senses, in particular to their whiskers, to gather the necessary sensory information and instruct navigation. Previous research has shown that rodents can process whisker input to guide mobility even prior to whisking onset by the end of the second postnatal week, however, when and how adaptive sensorimotor control of whisker position matures is still not known. Here, we addressed this question in rats longitudinally as animals searched for a stationary target in darkness. The results showed that juvenile rats perform object localization by controlling their body, but not whisker position, based on the expected location of the target. Adaptive, closed-loop, control of whisker position matures only after the third postnatal week. Computational modeling of the active whisking showed the emergence of the closed-loop control of whisker position and reactive retraction, i.e. whisker retraction that ensures the constancy of duration of tactile sampling, facilitate the maturation of sensorimotor exploration strategies during active sensing. These results argue that adaptive motor control of body and whiskers develop sequentially, and sensorimotor control of whisker position emerges later in postnatal development upon the maturation of intracortical sensorimotor circuits.

## Introduction

From single-cell organisms to mammalians, all animals employ adaptive mobility schemes for navigation (Hughes and Celikel, 2019). Actively whisking rodents, e.g. rats and mice, highly depend on their whiskers (vibrissae) to navigate their environments as they are nocturnal species living in a sub-terrestrial habitat in the wild. During active whisking, motor commands are iteratively updated based on prior sensory information, propagating through the sensory axis (Voigts et al., 2008, 2015; Azarfar et al., 2018a). The change in motor state, in return, regulates the pattern of whisking and control spatiotemporal integration of sensory information to enable tactile navigation (Celikel and Sakmann, 2007; Voigts et al., 2008; Lim and Celikel, 2019). This process takes about 90 ms and requires integration of the sensory information acquired over the last three whisk cycles while sensory information collected in the current whisk cycle is plausibly used as an error signal (Voigts et al., 2015). The end result of this computation is the protraction of whiskers to the position where the animal “thinks” the target object is located, rather than the actual position of the target (Voigts et al., 2015).

Motor dynamics of whisker positional control is not the only parameter that governs adaptive positional control during navigation. Body position and head angle are iteratively controlled to optimize the acquisition of sensory information. Locomotion and postural aspects of navigational sensorimotor computation (how the animal supports, moves, and controls particular body parts) develop alongside the maturation of relevant neural circuitry (Geisler et al., 1993; Clarac et al., 2004; Lelard et al., 2006), similar to whisking behavior (Landers and Philip Zeigler, 2006; Inan and Crair, 2007; Li and Crair, 2011; Erzurumlu and Gaspar, 2012; Grant et al., 2012a; Vitali and Jabaudon, 2014).

Coincidental to the development of cortical neural networks (Hensch et al., 1998), most behavioral components of navigation matures by the third postnatal week. Rat pups exhibit a flutter response to passive whisker contacts starting from P3 (Sokoloff et al., 2015). They can retract their whiskers starting from P4 and protract them P7 onwards (Landers and Philip Zeigler, 2006). Large amplitude active whisking emerges around the second postnatal week (Mosconi et al., 2010) even though animals can process passive whisker contacts to drive body mobility prior to the onset of active whisking (Clem et al., 2008). Thereafter, whisking gradually increases in frequency and amplitude, and reaches the adult form by the end of the third week (Landers and Philip Zeigler, 2006; Arakawa and Erzurumlu, 2015) as animals start whisking in a stimulus-dependent manner (Grant et al., 2012b). Complementary changes in the body positional control also appear during this period. Before P11, whisker movements are largely limited to unilateral retractions followed by head turns; between P11 and P13 bilateral whisking develops along with enhanced forward locomotion and improved control of the head position (Grant et al., 2012a). Contact-induced modulations of whisking symmetry, synchrony, and whisking amplitude modulation emerge shortly thereafter and continue to develop at least until ~P18 when adult-like locomotion patterns (Lim and Celikel, 2019), including rearing, develops (Grant et al., 2012a).

Considering that body and whisker positional control mature in parallel, it's tempting to speculate that sensorimotor computations that allow animals to adaptively control the position of their whiskers based on the recent sensory information also emerge by the end of the third week. Here, we experimentally tested this hypothesis using publicly available (Azarfar et al., 2018b) high-speed videography recordings of rats performing a tactile object localization task on the gap-crossing task (Hutson and Masterton, 1986; Celikel and Sakmann, 2007; Voigts et al., 2008, 2015; Pang et al., 2011; Juczewski et al., 2016; Miceli et al., 2017). Rats performed object localization first as a juvenile (P21) and later as adults (~P65). Quantitative analysis of whisking, body position, and tactile exploration showed (surprisingly) that juvenile animals fail to utilize sensory information to drive adaptive motor control of whisker position; they continue whisking without altering the whisk amplitude and set-points in the whisk cycle. Adult rats, however, perform the motor planning based on recent sensory information to actively control the midpoint and amplitude of whisking, as previously shown in mice (Voigts et al., 2015). Computational simulations and behavioral observations of the active and adaptive motor control of whisker position argue that the brain must utilize a neural code that allows reactive retraction of whiskers that keep the duration of contact roughly constant across touch events. We propose that the emergence of reactive whisker retraction along with the adaptive control of midpoint of whisk cycle (Voigts et al., 2015) are essential for the maturation of adaptive whisker positional control during navigation, which in turn (as we show) minimizes the amount of sensory information collected during goal-directed navigation.

## Materials and methods

All experimental procedures have been performed in accordance with the Dutch and European laws concerning animal welfare, and as per the guidelines for the care and use of laboratory animals, upon institutional ethical committee approval. The raw data have been previously made available as a part of the tactile navigation database (Azarfar et al., 2018b) and are distributed under a Creative Commons CC-BY license which permits unrestricted use, distribution, and reproduction of the data in any medium provided that the database (Azarfar et al., 2018b) is properly cited.

The spontaneous tactile object localization (Celikel and Sakmann, 2007) experiments were performed on 11 male Wistar rats (Harlan Laboratories, The Netherlands). Animal handling, behavioral observations and behavioral experiment paradigm were as described before (Azarfar et al., 2018b) and were comparable to the previous studies utilizing the same task (Celikel and Sakmann, 2007; Voigts et al., 2008, 2015; Pang et al., 2011; Juczewski et al., 2016; Miceli et al., 2017). In short, on this task animals shuttle between two elevated platforms with a variable distance between them as a camera placed over the gap between the two platforms is used to visualize the body and whisker position (Fig.1A). For the experiments analyzed herein, the data were collected using an AVT Pike (Allied Vision, Germany) camera at 220fps with a spatial resolution of 625 micron/pixel. All recordings were made in darkness. The gap between platforms was illuminated using a custom infrared (890 nm) backlight. Gap-distance was randomly selected from a normal distribution. The animals did not receive any reward for successful task execution. The animals had all of their whiskers intact until the last experiment for which all whiskers but one (either C2) was deprived under isoflurane anesthesia 24h before the behavioral experiments. To ensure that the data is not confounded by repetitive sensorimotor training and that it comes from a single day, we constrained the data to <3 trials/animal (N=30 per age condition). We analyzed only those trials where animals successfully located the target platform and crossed the gap to ensure that behavioral measures are not confounded by the trial outcome. Elevated platforms and the camera were mobilized by robotic actuators, thus animals were not exposed to human experimenters during behavioral observations.

**Figure 1.**
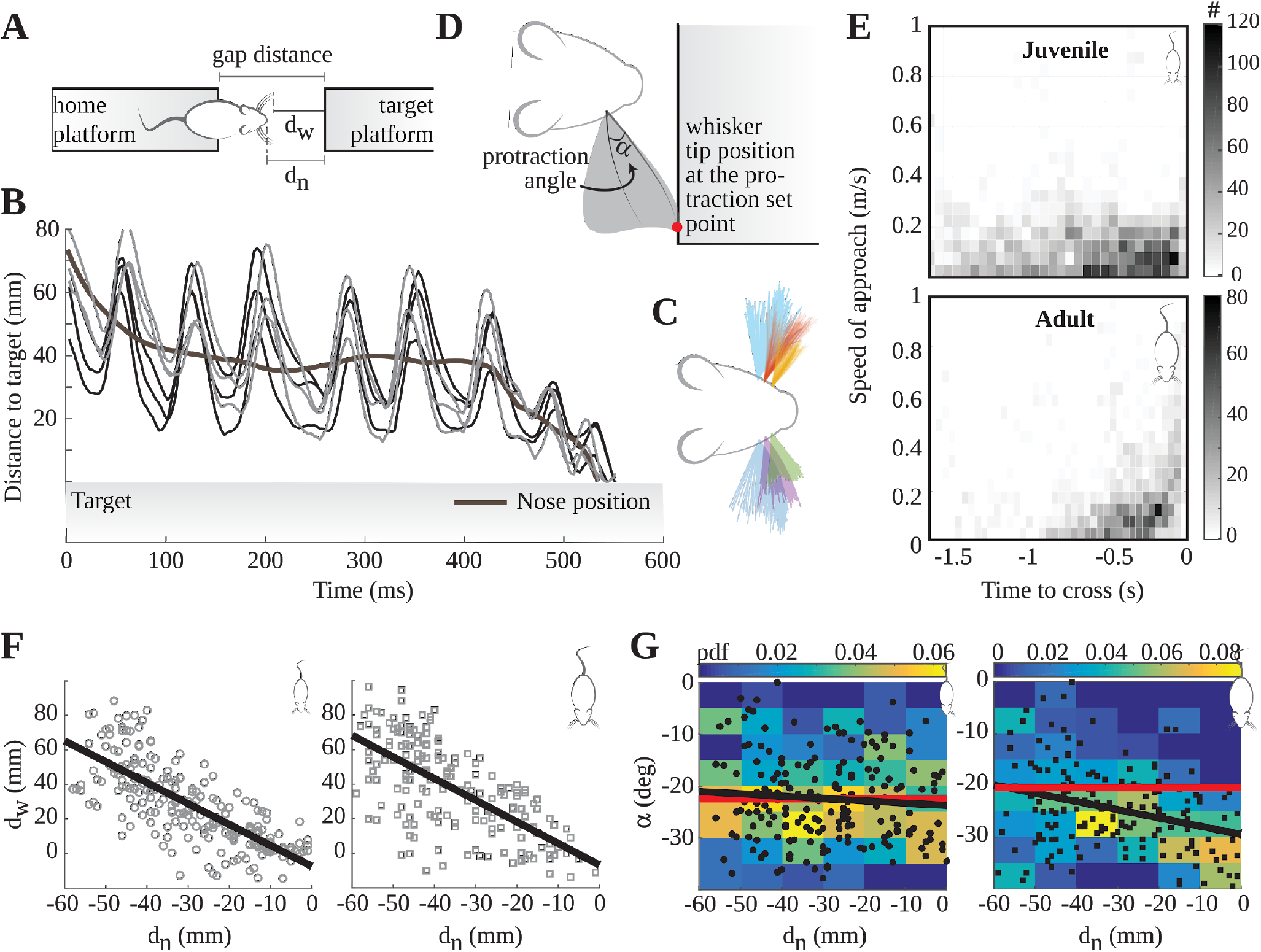
Motor strategies for tactile navigation. **(A)** The spontaneous tactile object localization training on the gap-crossing task. Animals shuttle between two elevated platforms separated with a gap in between. d_n_ : nose distance to the target, d_w_ whisker’s tip distance to the target. **(B)** An example of a representative dataset of target position, animal position, approach curve, and whisker tracking results. **(C)** Overlay of the tracked whiskers. Data from a single trial of a juvenile animal is shown. **(D)** Graphical representation of the protraction angle (**α**). Grey shaded area represents the whisker position during the course of a trial, normalized to the positional change in the body and angular change of the head. **(E)** The speed of locomotion during the last 1.5 seconds prior to gap-crossing. Top: juveniles (F = 1.36; p = 0.13 ANOVA); bottom: adults (F = 49; p <0.01 ANOVA). **(F)** Whisker tip distance to the target (d_W_) in relation to the body-to-target position. Left: Juveniles, r^2^ = .52, Nose distance effect: F = 4.82; p<0.001; Right: adults, r^2^ = .47, Nose distance effect: F = 2.92; p<0.001. **(G)** The most protracted angle at each whisk cycle in relation to the setpoint versus the nose distance d_n_ from the target. Left: Juveniles, Fit slope= 0.03 deg/mm, Nose distance effect: F = 0.85; p=0.09. Right: Adults (Fit slope= - 0.17 deg/mm, Nose distance effect: F = 1.59; p<0.01). Black lines are linear fits to the data, red lines are “null correlation” fits, i.e. the trend expected if the d_n_ and **α** were not correlated.

### Analysis of tactile exploration

All images were analyzed in MATLAB. A background image was selected before the animal entered the field of view and was subtracted from all subsequent frames. Platform edges were detected in the background picture as two transition points on a back-lit background. To determine the head angle, a triangle was created between the two ears and nose. Deviation of the nose from the head center (between the ears) was used to calculate the head angle. Whisker contacts with the target platform and a select subset of caudal whiskers (C1-4) were tracked manually (N=6 whiskers/animal; N=3 whiskers/snout). The angular disposition of the base of the whisker was analyzed temporally and spatially (in respect to body-to-target position, i.e. nose to edge of the target platform). The speed with which animals approached the platform was quantified using the time derivative of the distance between the animal’s nose and the edge of the target platform. The frequency of whisking was calculated using Fast-Fourier Transform (FFT) and from the cycle duration (i.e. the time it takes for the whisker to travel between two protraction set-point). Gap-crossing was defined as the moment the animal’s nose has passed the platform edge by more than 6 mm. Videos were then grouped by age and gap distance. Only those trials where the tactile target was at a “whisker distance”, i.e. 14.5+ cm for adults and 9+ cm for juvenile, were used for further analysis. These distances were selected to ensure that only tactile exploration of whisker touch was quantified, as animals also use the touch receptors around their nose in shorter gap-distances (Celikel and Sakmann, 2007; Voigts et al., 2008, 2015; Pang et al., 2011; Juczewski et al., 2016; Miceli et al., 2017). Whisker protraction was quantified in respect to the midpoint of the whisk space which was defined by the most protracted and retracted positions, normalized to the nose position, during the tactile exploration epoch.

## Results

Quantitative analysis of the interplay between motor control and sensory acquisition strategies in freely exploring animals will help to unravel the principles of sensorimotor computation for navigation. Here, by observing juvenile (P21) and adult (~P65) rats locate stationary targets we describe the development of adaptive motor control in the whisker system.

### Motor strategies for tactile navigation across development

On the gap-crossing task, animals navigate between two elevated platforms with a variable distance between them in darkness (Celikel and Sakmann, 2007; Voigts et al., 2008, 2015; Pang et al., 2011; Juczewski et al., 2016; Miceli et al., 2017). They extend their body over the gap to reach the tactile target using their whiskers (Fig.1A). To analyze animals’ locomotive strategies, we studied the speed at which they approach the target in the rostrocaudal orientation. This maneuver is a goal-directed behavior that minimizes the relative distance to the target, maximizing the likelihood of the first contact with the target.

Spatially resolved speed of approach to the target showed that juvenile animals do not vary approach speed (Fig.1E, top) although adult animals approximately double it every 100 ms during the last 400 ms prior to gap-cross (Fig.1E, bottom). Even though this approach profile could be interpreted as juvenile rats navigate towards the target without adapting their motor strategy, increased mobility could also be a by-product of non-motor variables (e.g. change in exploration vs exploitation strategies during development).

Whisking rodents control the angular position of their whiskers by independently controlling whisker protraction and retraction during tactile navigation (Sachdev et al., 2002; Voigts et al., 2008, 2015; Sofroniew and Svoboda, 2015). To decode the contribution of whisking motor pattern modulations on the sensorimotor computation, we quantified the angular disposition at the base of the whisker during tactile exploration. Analysis of whisker tip position with respect to the target revealed that the whisker tip position converges on the tactile target in both juvenile and adult animals as the animal approaches the target (Fig.1F), suggesting that juveniles can perform adaptive positional control to locate stationary targets in their immediate environment. These results, however, do not reveal whether they also actively control their whisker position during tactile exploration. Thus we next quantified whisking patterns. Maximally protracted and retracted whisker positions, normalized to the nose position, were used to visualize the “whisk space” and the middle of this range was set as zero degrees set-point (also called midpoint, or whisker position at rest (Voigts et al., 2008, 2015)). The protraction angle (**α**; Fig.1D) was calculated with respect to the zero degrees setpoint. Quantification of the change in the protraction angle (Fig.1G) showed that adult rats position their whiskers more rostrally (bring them forward) as they approach the target, as previously shown in mice (Voigts et al., 2015). Juvenile animals, on the other hand, do not regulate their whisker protraction independent from their relative distance to the target and sensory information acquired in the previous whisk cycles. These results argue that adaptive whisker positional control matures after P21.

### Frequency and amplitude modulation across development

During active sensing, sensory exploration is contextually modified in a task and goal-specific manner in part by whisking frequency modulation (Carvell and Simons, 1990, 1995; Voigts et al., 2008, 2015; Mitchinson et al., 2011; Zuo et al., 2011). Berg and Kleinfeld (Berg and Kleinfeld, 2003) reported an increase in the whisking frequency range when animals change their whisking mode from ‘exploratory’ (range 5–15 Hz) to ‘foveal’ whisking (15–25 Hz) in search for a reward. However, it is not yet known whether tactile sampling strategies adapt after the contact with the target. Considering that the higher the whisking frequency is, the shorter the duration of tactile exploration at a given whisk cycle, reduction in whisk frequency during tactile exploration will maximize the contact duration unless whisker retraction is actively modulated (see below).

#### Adaptive frequency modulation develops postnatally after P21

Whisking frequency distribution mostly lies between 5 and 20 Hz for both juveniles and adults during stationary target localization, independent from the animals’ location with respect to the target (Fig.2A). In juvenile animals tactile exploration does not change the whisking frequency (Fig.2B, top), although adult rats reduce the frequency of whisking upon the first contact with the target (Fig.2B, bottom). Considering that protraction and retraction amplitudes during free-whisking are comparable across animals (Fig.2C), contact-induced change in whisking frequency is likely to be mediated neuronally, by controlling the phasic regulation of whisking, for example by a delay line. In agreement with this observation, the comparison of the change in peak-to-peak amplitudes of whisking showed that adult animals adapt their whisk amplitude by a factor of three as they approach the tactile target (Fig.2D).

**Figure 2.**
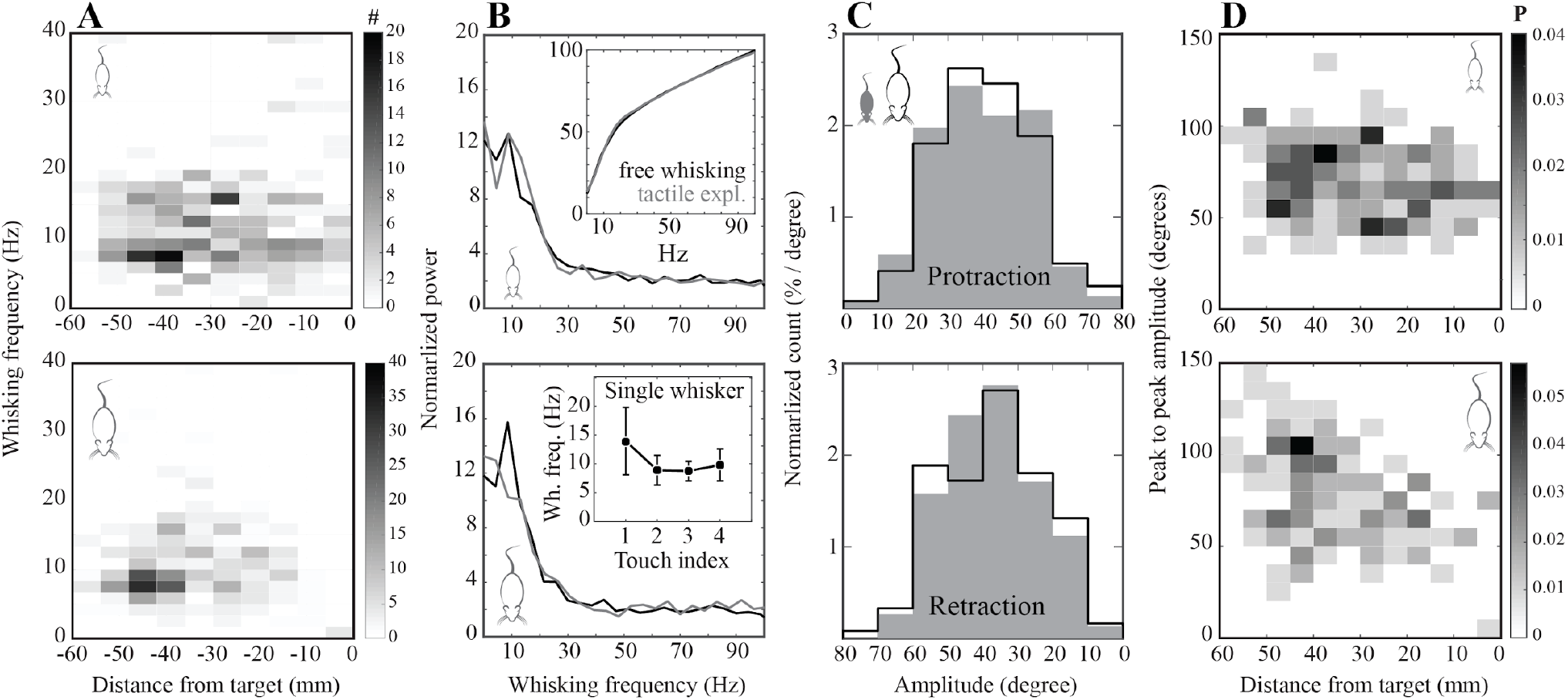
Frequency and amplitude modulation in juvenile and adult rats. **(A)** Binned histogram of whisking frequency in relation to the body-to-target distance (5 mm/bin; Top: Juveniles, F=1.36, p=0.07, Two-way ANOVA; Bottom: Adults, F=12; p=0.01, Two-way ANOVA). **(B)** Normalized power of frequency range in free whisking versus during tactile exploration. Top: Juveniles (whisking mode effect: F=0.75, p=0.22, ANOVA); inset shows the cumulative frequency distributions (whisking mode effect: F=0.57, p=0.79, ANOVA). Bottom: Adults (whisking mode effect: F=13.83, p<0.01, ANOVA); inset: whisking frequency binned by touch (data from rats with single whisker, N=5 rats; N=30 trials). (**C)** Normalized distribution of protraction and retraction angles in each whisk cycle (Top -- Protraction: developmental effect, F=1.78, p=0.27, ANOVA. Bottom -- Retraction: developmental effect, F=3.43; p=0.18, ANOVA).

### Sensorimotor adaptation and sensory acquisition across development

Analysis of the tactile exploration duration showed that adult rats spend significantly less time collecting sensory information compared to juveniles prior to successful object localization (Fig. 3A); even though the duration of single-touch events is statistically comparable between the groups (Fig. 3B). Accordingly, the difference in sensory exploration is due to the increased number of whisker contacts in juvenile animals (Fig. 3C). Because juveniles do not modulate zero degrees set-point (midpoint) of whisking and fail to adapt the amplitude of whisking based on their relative distance to the target (see above), most of their whisker contacts should be made later during tactile exploration.

**Figure 3.**
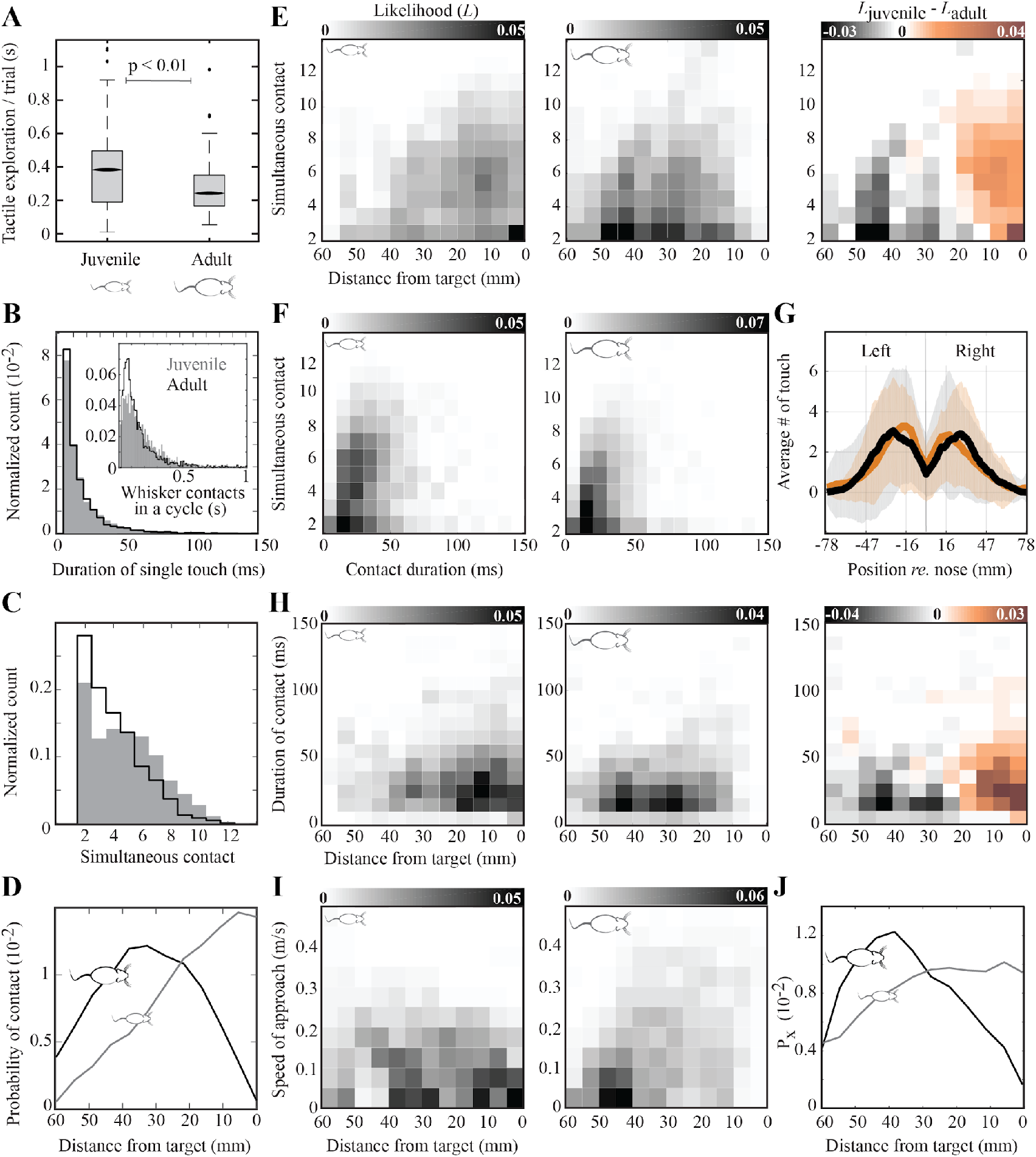
Statistics of whisker guided tactile exploration across development. **(A)** Adult rats spend significantly less time exploring the gap compared to juveniles prior to successful object localization (developmental effect: F=87; p<0.01). **(B)** Distribution of the duration of single-touch events, Grey: juveniles, white: adults (developmental effect: F = 0.74; p=0.49). The inner figure shows the total duration of contact events in a whisk cycle across all the whiskers (developmental effect: F=2.45; p=0.21). **(C)** Distribution of the number of whiskers used during tactile navigation (developmental effect: F =27; p<0.01). **(D)** The spatial probability distribution of contacts with respect to the relative distance to the target (developmental effect: F=251; p<0.01). **(E)** Spatial distribution of likelihood of the simultaneous number of contacts during tactile navigation (developmental effect (observed variable distance): F=221; p<0.01). **(F)** Histograms of average contact duration grouped by the number of whiskers simultaneously in contact with the target (developmental effect (observed variable contact duration): F=73; p<0.01). **(G)** Normalized distribution of contacts in the horizontal plane (developmental effect: F=0.364; p=0.546). **(H)** Histograms of contact duration grouped by the relative distance to the target (developmental effect (observed variable distance): F =198; p<0.01). **(I)** Distribution of speed of approach by relative distances to target (developmental effect (observed variable distance): F=369; p<0.01). **(J)** Normalized distribution of time spent with respect to relative distance to the target (F =183; p<0.01). All statistical comparisons are made using (two-way) ANOVA.

#### Juvenile rats collect most of the sensory information closer to the target, recruiting a larger number of whiskers for tactile exploration, in comparison to adults

Juvenile and adult rats employ two different strategies to harvest sensory information. Adults make most of their contacts with the target while keeping their body within 2-5 cm of the target; in juveniles, the likelihood of sensory exploration increase as they approach the target (Fig. 3D). Deconvolution of the number of simultaneous whisker contacts with the relative distance to the target (Fig. 3E) showed that adult rats recruit a relatively small number of whiskers to simultaneously collect sensory information while juvenile animals engage larger number whiskers for exploration, in particular when they are within ~2 cm of the tactile target (Fig.3E, right). Because the contact duration and number of whiskers that are simultaneously in contact with the target is positively correlated in juveniles, but not in adult animals (Fig. 3F), juvenile animals effectively collect disproportionate amount of sensory information for tactile object localization (Fig. 3A), due to prolonged contact duration (Fig.3H), reduced speed of mobility (Fig.3I) and increased time spent (Fig.3K) as they approached the platform.

### An *in silico* model of active whisking

The results thus far show that juvenile animals do not utilize the sensory information to adapt their whisking patterns during tactile navigation. To systematically address how adaptive motor control can be implemented without considering the prior sensory information, we used an *in silico* model of whisking (Whisking *in silico*; github.com/DepartmentofNeurophysiology/Whisking-in-silico). The whisking module of the model is based on a pair of coupled oscillators driving protraction and retraction, functionally mimicking the central pattern generators in the brainstem (Cramer and Keller, 2006; Moore et al., 2014). As in the biological circuits of sensorimotor control, these oscillators receive top-down hierarchical input that could modulate the whisk cycle (Cramer and Keller, 2006; Moore et al., 2014). The model integrates body locomotion and whisking motor control, thus can decouple the two active motor control processes to address how adaptive changes in body positional control relate to adaptive whisking. Sensory feedback, e.g. duration of contact, can be integrated to establish closed-loop sensorimotor control. Using this model, non-adaptive (uniform) whisking, as observed in juvenile animals, can be simulated by driving the oscillators as an open-loop, i.e. the whisking frequency, protraction and retraction onsets are not modulated by sensory information but intrinsically regulated.

#### Open-loop whisking replicates the pattern of sensorimotor exploration observed in juveniles

During open-loop (uniform) whisking, the whisker protraction amplitude (Fig.4B) and protraction set-point (Fig.4C) do not change based on the history of tactile exploration and relative distance of the animal to the target, similar to our behavioral observations in juvenile rats (Figs.1,2). Adult animals, on the other hand, employ adaptive whisking and systematically protract their whiskers further as they get closer to the target. *In silico*, such a control system requires closed-loop integration of sensory and motor circuits which is implemented through a recursive increase in protraction angle and a decrease in whisking amplitude upon the emergence of novel sensory information. This implementation successfully replicates the observed properties of adaptive motor control in adult animals as adaptive whisking *in silico* results in forward placement of the whiskers, increased whisker protraction amplitude, reduced amplitude upon contact and frequency of whisking (Fig.4B-F).

**Figure 4.**
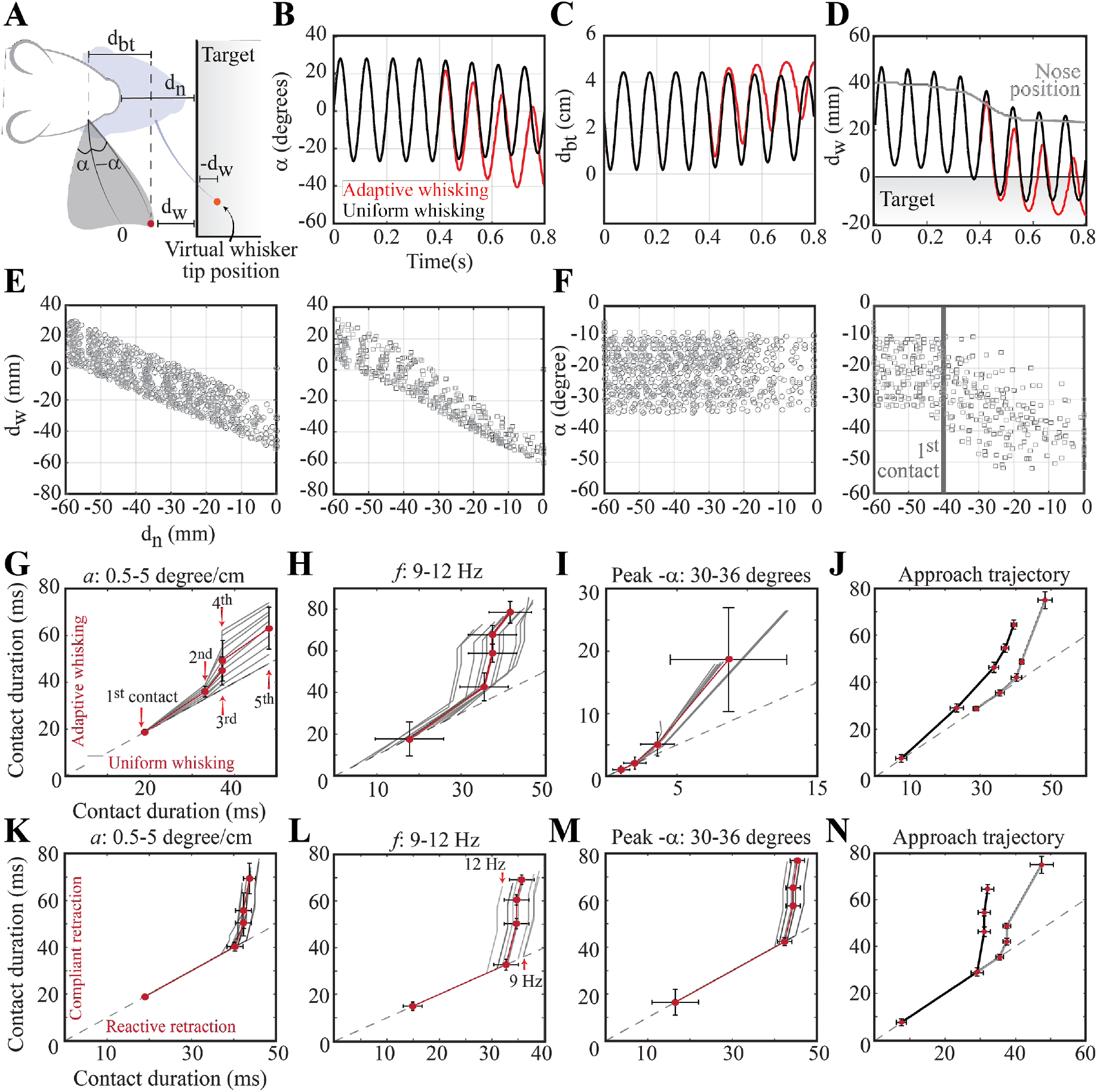
Whisking *in silico*. **(A)** Graphical representation of the variables used for visualization: protraction angle (−**α**), retraction angle (**α**), relative distance to target with respect to nose (d_n_), relative distance of the whisker tip to target (d_w_,−d_w_), distance between whisker’s base on the snout and whisker’s tip (d_bt_). The color code represents the relative distance to the target. **(B)** Uniform (open-loop, shown in black) versus adaptive (close-loop, red) whisking as observed in juvenile and adult animals, respectively. Adaptive whisking is characterized by a reduction in whisk amplitude, forward placement of whisker, increased protraction amplitude. During uniform whisking animals whisk at a constant frequency and amplitude. **(C)** Evolution of d_bt_, i.e. how far the animal is protracting its whiskers, during adaptive whisking (red) versus uniform whisking (black). **(D)** Change in the relative position of the whisker tip with respect to target during tactile exploration. Nose position is based on behavioral experiments; an approach trajectory from a randomly selected trial is shown. **(E)** Simulated tip distance to target in relation to the body-to-target position. Left: uniform whisking, right: adaptive whisking. **(F)** Adaptive motor control of whisker angular protraction. Left: uniform whisking, right: adaptive whisking. **(G-N)** Contact duration in adaptive versus uniform whisking. The dashed line shows the result for uniform whisking in each condition. **(G)** Sensory driven adaptive gradual increase of mid-point (a) increases the contact duration. Labels are the order of whisker contact during tactile exploration. **(H)** The relationship between the whisk frequency and contact duration during adaptive vs uniform whisking. **(I)** The interaction between the maximum protraction angle and the contact duration. **(J)** The role of body positional control for contact duration. The approach pattern entrained by juvenile rats (gray) results in higher contact duration in both open-loop and close-loop whisking. **(K-N)** Change in contact duration during compliant versus reactive retraction. The dashed line shows the result for compliant retraction in each condition. The contribution of the zero-degrees set-point (mid-point) adaptation **(K)**, whisking frequency **(L)**, maximum protraction angles **(M)** to the modulation of contact duration. **(N)** The role of approach trajectory in juveniles (gray) and adults (black) in shaping the contact duration during compliant versus reactive retraction. In G-N each red dot represents a single whisker contact; the red line is the average change in contact duration across the parameter space noted on each figurine.

#### The emergence of close-loop whisking for adaptive control of whisker position increases contact duration

Considering that close-loop whisking accurately replicates the change in body and whisker positional control across development, it is plausible that the transition from open-loop to close-loop whisking can explain the differences in sensory exploration between juveniles and adults. Therefore, we compared contact (touch) durations across active whisking strategies *in silico* (Fig4.G-J). The body locomotion was the same for the first three experiments (Fig.4C-E) and based on the approach trajectory (i.e. nose distance to the target) shown in Figure 4D.

Whisking *in silico* provides means to decouple motor control variables and study their contribution to sensory exploration individually. Because juvenile animals fail to adapt the zero-degrees set-point (midpoint) of whisking, we first addressed the contribution of midpoint adaptation to contact duration (Fig.4G). The results showed that as the midpoint adaptation increase in amplitude (similar to observed in adult animals), the contact duration is gradually prolonged. Although the contact duration of the first touch was comparable across open-loop (uniform) and close-loop (adaptive) whisking, with successive whisker contacts the difference increased in favor of the adaptive whisking (Fig.4G). Contact duration during open-loop whisking changed only minimally independent from the change in midpoint adaptation (Fig.4G), whisking frequency (Fig.4H) and whisk protraction amplitude (Fig.4I), suggesting that it is primarily modulated by the animal’s relative body position to target. To test this hypothesis, we ran additional simulations with the same whisking parameters (Fig4G-I) but using the approach trajectories of juvenile or adult rats (Fig.4J). Independent from the active whisking strategy *in silico*, i.e. open-loop *versus* closed-loop whisking, the contact duration was longer with juveniles’ approach trajectory (Fig.4J), confirming that body positional control has a pronounced contribution to contact duration in juveniles. The results of *in silico* whisking outlined thus far show that the switch from uniform to adaptive whisking replicates the observed developmental changes in whisker positional control during tactile navigation, however it prolongs the duration of individual contacts, thus failing to reproduce the constancy of contact duration across juveniles and adults (Fig.3B).

#### Reactive retraction shortens the contact duration, counteracting the prolonged whisker contacts observed upon the emergence of adaptive whisking

From the motor control perspective, the contact duration might be regulated by “reactive retraction” where sensory information is used to modulate the contact duration during closed-loop sensorimotor control. Alternatively, if the sensory input does not have any influence on the retraction of whisking, the retraction would be compliant to the protraction (“compliant retraction”) as they are driven by a dual-phase central pattern generator without any need for sensory input (Berg and Kleinfeld, 2003). Compliant retraction, however, results in prolonged contact duration *in silico* (Fig.4K-N) while any change observed during the reactive reaction is a result of the change in approach trajectory (Fig.4B), thus due to body positional control (Fig.4N).

To experimentally test the prediction of the *in silico* model, and to address which of the two models of retraction is employed by animals, we quantified the touch duration and duty cycle across consecutive touches (unilateral C2 whisker; Ns=5 animals, 30 trials; Fig.5). The results showed that touch duration increases after the first contact as the animals get closer to the target and remains constant thereafter (Fig.5A), while the duty cycle remains constant across all touch events (Fig.5B); in agreement with previous findings (Deutsch et al., 2012). Unlike these behavioral observations, in simulated whisking (Fig.4K-N) whisker contact duration does not reach an asymptote. It is because in the model the retraction time is based on the sensory information in the previous cycle and a reference motor copy of the current whisk cycle. Thus, body dispositions since the last whisker contact and the sensory information in the current whisk cycle are not included, although, as behavioral recordings show, it is utilized by animals for the reactive retraction of whiskers. These results suggest that tactile (whisker) information driven closed-loop control of whisker position during development not only modulates the amplitude of whisking but also enables “reactive retraction” to ensure constancy of whisker contact duration across touches as sensorimotor strategies for tactile navigation mature.

**Figure 5.**
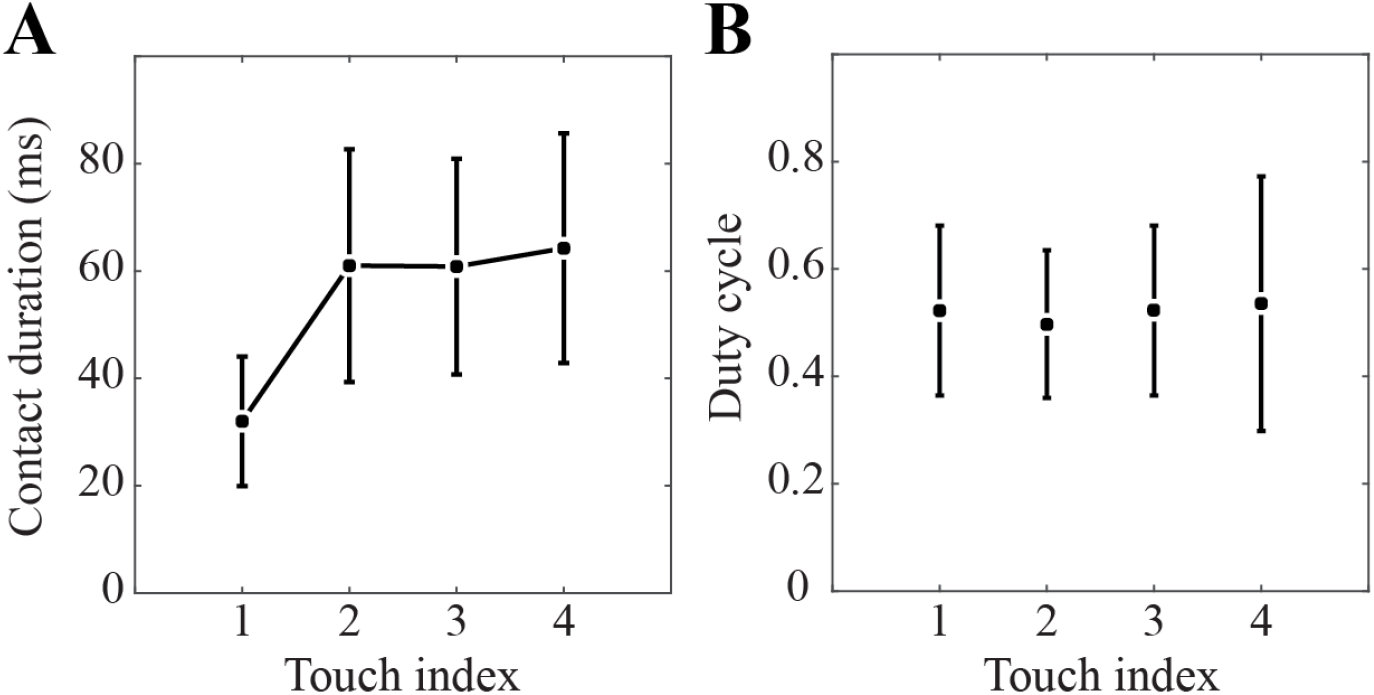
Sequential change in contact (touch) duration and duty cycle across whiskers contacts. Data from adult animals with a single spared whisker (unilateral C2; N=5 animals and 30 trials). **(A)** Contact duration as a function of the touch index. Note that after the first contact, the contact duration is stabilized. **(B)** The duty cycle of whisking, i.e. duration of contact with respect to the duration of a whisk, across the different touch events.

## Discussion

In order to unravel the interplay between the motor control and sensory acquisition strategies, and to address the developmental changes in sensorimotor control, here we studied the whisker and body positional control during tactile navigation in juvenile (P21) and adult (~P65) rats. Quantitative analysis of behavior showed that both groups of animals successfully locate tactile targets although they do so using different sensorimotor strategies (Fig.6). Juvenile animals control their body position to maximize the sensory information acquisition as they continue whisking uniformly, independent from their distance to the tactile target (Fig.6A,B). Adults, on the other hand, control where and how they whisk based on the sensory information they acquire during tactile exploration; they protract their whiskers further, they reduce the whisker protraction amplitude and perform reactive whisker retraction (Fig.6C). As we computationally showed, the change in sensory and motor exploration patterns during development can be explained by the emergence of closed-loop sensorimotor control, through which the tactile information collected during the course of exploration is used to plan where to whisk next. These simulations predicted that adaptive changes in whisker positional control have to be accompanied by the recruitment of reactive retraction of whiskers to stabilize contact duration, which we confirmed behaviorally (Fig.5).

**Figure 6.**
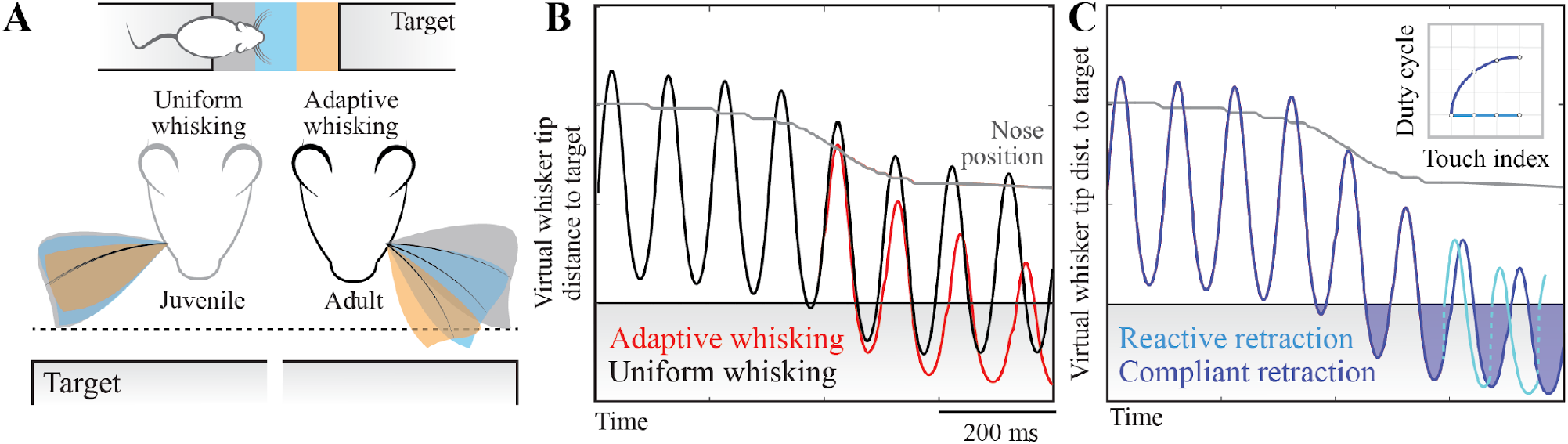
Developmental changes in sensorimotor navigation during goal-directed tactile exploration. **(A)** Uniform versus adaptive whisking as observed in juvenile and adult animals, respectively. The color code represents the relative distance to the target (top) and the motor control phenotype at the corresponding distance (middle). Juvenile animals whisk uniformly at a constant frequency and amplitude independent from their distance to the target and the sensory information collected during exploration. Adaptive whisking in adults is characterized by a gradual reduction in whisk amplitude, forward placement of whisker and increased protraction amplitude as the animal approaches the target. See Figs.1–3 for the behavioral data, see Fig.4 for how the change in motor control variables alters the acquisition of sensory information. **(B)** The emergence of adaptive whisking results in increased contact duration (sensory exploration) with the target. **(C)** Emergence of reactive retraction counteracts the increased exploration as the relative duration contact, i.e. duty cycle, is kept constant across subsequent whisks during tactile exploration (see Figure 5 for behavioral data).

A direct outcome of reactive retraction is that the whisking duty-cycle is kept constant (Fig. 5). We observed that the touch duration increases from the first touch to the second as animals approach the target and remain constant during the rest of the tactile exploration. These results are in agreement with the reduction in the frequency of contact after the first touch (Fig.2) and provide independent support for the closed-loop control of whisker positional control during tactile navigation.

### The choice of developmental age

For the experiments with juveniles, we focused on postnatal (P)21 primarily because, at this age, the whisker system of rats is believed to be functionally (see introduction) and anatomically mature. The barrels begin to emerge at E9 (prenatal) and continue to form characteristics of its mature network until P21 (Hensch et al., 1998). Patterning and plasticity of the barrel cortex, the targeting of ventrobasal thalamic axons, the formation and functional maturation of synapses during the initial critical developmental periods are extensively investigated (Inan and Crair, 2007; Li and Crair, 2011; Erzurumlu and Gaspar, 2012; Vitali and Jabaudon, 2014). Initially, through corticogenesis, the intrinsic genetic programs determine cortical arealization. Thereafter, thalamocortical axons (TCA) innervate the cortical plate (McConnell et al., 1994; Del Río et al., 2000), barrels are formed around P3 and mature by P7 (Persico et al., 2001; Fox, 2002). Around P14, TCA projections innervate the barrel cortex (Bender et al., 2003) and central pattern generators in the brainstem start controlling whisker motion (Landers and Philip Zeigler, 2006). In the third postnatal week, the pattern of GABA immunoreactivity, the receptive field of excitatory neurons and their axodendritic projections reach maturity (Del Rio et al., 1992; Bender et al., 2003). Considering that both body positional control and the whisker motion mature during this period (see introduction), this age group provides a suitable time course to address the principles of adaptive sensorimotor control in juveniles.

### The function of adaptive whisking

Adaptive sensorimotor computation, including adaptive whisking, has been proposed to maximize the task-related sensory information and minimize energy consumption (Nelson and MacIver, 2006; Hofmann et al., 2013). In behavioral tasks like stationary object-localization (Celikel and Sakmann, 2007) adaptive whisking might also minimize the redundancy in the incoming sensory information. With a stationary tactile target that has a uniform height, surface structure, and stiffness, and explored in darkness, egocentric and allocentric sensory information available to the animal across different whisker is mostly redundant; therefore strategies described in more complex (navigational) tasks (e.g. (Shimshek et al., 2006; Celikel et al., 2007; Corsini et al., 2009; Freudenberg et al., 2013, 2016; Seib et al., 2013; Pezzulo et al., 2014; Spiers and Gilbert, 2015; Heckman et al., 2016, 2017; Behrens et al., 2018; Górska et al., 2018) are likely to be inconsequential. Considering that animals can perform the task equally well even with a single intact whisker (Celikel and Sakmann, 2007) and that the emergence of adaptive-whisking results in recruitment of smaller number of whiskers during tactile object localization (Fig.3C), it is plausible that adaptive whisking could be a strategy to reduce the redundancy in sensory information even before a sensory neuron encodes the information available in the periphery. Besides redundancy reduction, adaptive motor control governs the pattern of whisker movement, which dictates the sensory signal transmitted along each whisker and the exploratory field of each whisker (Azarfar et al., 2018a). This could potentially influence the sensory information available to the brain to solve the control task.

### Outlook

Sensorimotor systems utilize probabilistic models to predict unobserved variables (Wolpert and Flanagan, 2016). The brain generates a prior hypothesis based on its internal models of the external world and combines them with acquired sensory information to form a posteriori hypothesis of the target location (Wolpert et al., 2011; Voigts et al., 2015). As the animal approaches the target during tactile navigation, acquired sensory information in egocentric coordinates, modulate the posterior hypothesis. Using learned forward models the posterior could be translated into motor action to drive adaptive sensing. Updating the internal models could result in learning these forward models using the sensory error signals to learn the inverse models (Taylor et al., 2014). Our results suggest that acquisition of forward models, which is likely to be experience-dependent, happens after P21 upon the maturation of intracortical sensorimotor circuits. Longitudinal chronic recordings from sensory and motor cortices starting from the last week of the first postnatal month will shed light on the developmental (including experience-dependent) and neuronal mechanisms of sensorimotor computation. Combined with the systematic molecular, structural and functional reconstruction (see e.g. (Meyer et al., 2010, 2013; Narayanan et al., 2015; Huang et al., 2016; Kole et al., 2017b, 2017a, 2018; da Silva Lantyer et al., 2018; Azarfar et al., 2019; Kole and Celikel, 2019)) of these circuits, a comprehensive multi-scale computational map of sensorimotor computation is evolving.

